# Bean yellow mosaic virus infecting broad bean in the green belt of Córdoba, Argentina

**DOI:** 10.1101/606384

**Authors:** Patricia Rodríguez Pardina, Claudia Nome, Pablo Reyna, Nacira Muñoz, Evangelina Argüello Caro, Andrés Luque, Humberto Julio Debat

## Abstract

Broad bean (*Vicia faba* L) is the fourth most important pulse crop in the world. In Argentina, broad bean production was of 1,841 hectares and 16,500 tons during the 2017 growing season. Broad bean is commonly used in rotations; especially by farmers located in “green belts” that are peri-urban areas surrounding large cities that include horticultural family farms. Plants showing marked foliar mosaic symptoms, typical of viral infection, were collected during the 2015 growing season in the green belt of Córdoba city, Argentina. Preparations of symptomatic tissues were mechanically inoculated onto healthy broad bean plants in the greenhouse, which developed symptoms similar to those observed in the field. In addition, symptomatic samples were positive when tested by indirect ELISA with the *anti*-*potyvirus* group monoclonal antibody. Further, flexuous filamentous particles typical of potyviruses were observed under the electronic microscope on dip preparations. Lastly, total RNA was extracted from a symptomatic leaf and high-throughput sequenced, which allowed the assembly of a single virus sequence corresponding to a new highly divergent strain of *Bean yellow mosaic virus* (BYMV). Phylogenetic insights clustered this Argentinean broad bean isolate (BYMV-ARGbb) within group IX of BYMV. Given the economical importance of this virus and its associated disease, the results presented here are a pivotal first step oriented to explore the eventual incidence and epidemiological parameters of BYMV in broad bean in Argentina.

## Introduction

Broad bean (*Vicia faba* L) is one of the oldest cultivated vegetables. This species was originated in the old world (Afghanistan, Western Asia and the surroundings of the Himalayas), and it is the fourth most important pulse crop in the world with a global production of 2.5 million hectares and 4.8 million tons during the 2017 growing season (FAOSTAT 2017). The main producer of this legume is China, where it has multiple uses, such as consumption as dry grain, green vegetable or processed food, since it is a cheap source of high quality protein (Makkouk et al. 2003; Muñoz et al. 2017). Additionally, a distinctive characteristic of this crop is its ability to associate with atmospheric nitrogen-fixing bacteria in a symbiotic relationship. Therefore it is used in rotation systems since this interaction is the most inexpensive and environmentally friendly nitrogen source for crop production (Graham and Vance 2000).

The information about broad bean in Argentina is scarce, nonetheless according to FAOSTAT (2017) estimations, the nationwide production was of 1,841 hectares and 16,500 tons during the 2017 growing season. Broad bean is an autumn-winter crop and the mainly planted variety “Agua Dulce” is commonly used in rotations, especially by small family farmers in transition to agroecological systems (Goites 2008). These small family farms are located mainly in “green belts,” that are defined as peri-urban areas surrounding large cities that include horticultural family farms and eventually small sized business farms whose production is intended especially for fresh vegetables (Paladino et al. 2018).

Fungal, bacterial and viral caused diseases have been a serious limitation for broad bean production. The most important fungal diseases affecting broad been are: chocolate spot (*Botrytis fabae* and *B. cinerea*), rust (*Uromyces viciae fabae*), black root rot (*Thielaviopsis basicola*), stem rots (*Sclerotiniatrifoliorum, S. sclerotiorum*), root rots and damping-off (*Rhizoctonia* spp.), downy mildew (*Pernospora viciae*), pre-emergence damping-off (*Pythium* spp.), leaf and pod spots or blight (*Ascochyta fabae*), and foot rots (*Fusarium* spp.). Further, the most important viruses reported to infect broad bean worldwide include members of the *Bean leafroll virus, Bean yellow mosaic virus, Beet western yellows virus, Broad bean mottle virus* and *Faba bean necrotic yellows virus* species (Singh et al. 2012). Plants showing marked foliar mosaic symptoms, typical of viral infection, were collected during the 2015 growing season in the green belt of Córdoba city, Argentina; this work was aimed to identify the causal agent of the infected samples

## Materials and methods

Samples with mosaic symptoms were mechanically inoculated onto healthy broad bean plants, using 0.01M sodium phosphate buffer pH 7, 0.1% Na_2_SO_3_, plus silicon carbide (600 mesh). The plants were transferred into a greenhouse (20–24°C) and observed daily until symptoms development.

For Electron microscopy observation leaf dips were prepared with symptomatic leaves, ground in PBS pH 7 + 0.01% (w/v) sodium sulphite (Na_2_SO_3_). A drop of this extract was transferred to carbon-coated Formvar grids for seven min. The grids were then washed with distillate water, negatively stained with 2% uranyl acetate and examined with a JEM EXII transmission electronic microscope (Jeol, Tokio, Japan).

The infected plants were also tested by DAS-ELISA or PTA (Clark and Adams 1977; Mowat et al. 1987) with antisera specific for: *Alfalfa mosaic virus* (AMV), *Cucumber mosaic virus* (CMV), *Tomato spotted wilt virus* (ToSWV), *Tobacco streak virus* (TSV) and *Potyvirus*. The reactions were quantified after 60 min incubation at 410 nm (OD410) using a Sinergy H1 microplate reader (BioTek, USA). Samples were considered positive when OD410 values were higher than the mean of the healthy controls plus three times the standard deviations (six healthy controls per plate were used).

For molecular characterization total RNA was extracted from a symptomatic leaf using RNeasy Plant Mini Kit (Qiagen, Hilden, Germany) and 200 µl (129 ng/µl), were submitted to INTA Castelar for a next-generation sequencing (NGS). The samples were assessed by Bioanalyzer (Agilent, inc) and prepared for sequencing with a Miseq System (Illumina, inc) equipment. The obtained reads were processed and analysed for virus discovery as previously reported (Debat 2017). In brief, raw reads were filtered and trimmed with the Trimmomatic v0.38 tool as implemented in http://www.usadellab.org/cms/?page=trimmomatic and assembled *de novo* with standard parameters using Trinity v2.8.3 available at https://github.com/trinityrnaseq/trinityrnaseq/releases. The obtained transcriptome was subjected to bulk BlastX searches (E-value < 1e-5) against refseq virus proteins available at ftp://ftp.ncbi.nlm.nih.gov/refseq/release/viral/viral.2.protein.faa.gz. Virus hits were curated and reassembled as described in Debat and Bejerman (2019). Annotation was done with standard tools such as the ORFfinder https://www.ncbi.nlm.nih.gov/orffinder/ NCBI-CDD https://www.ncbi.nlm.nih.gov/Structure/cdd/wrpsb.cgi and HHPred https://toolkit.tuebingen.mpg.de/#/tools/hhpred. Detected domains, cleavage sites and sequence similarity were explored by hand using multiple sequence alignments (MSA) of predicted gene products using MAFFT v7 with “auto-select” algorithm parameter available at https://mafft.cbrc.jp/alignment/software/. In addition, sequences were compared with those of other viruses available in GenBank (https://www.ncbi.nlm.nih.gov/genbank/, Table 1). Maximum Likelihood (ML) phylogenetic trees were constructed with MEGA 5.2 program (Tamura et al. 2011), using the G nt substitution model and 1000 bootstrap iterations (CP protein) or MAFFT v7 with BLOSUM64 scoring matrix for amino acids and E-INS-i iterative refinement method (PP). Alignments were trimmed using GBlocks tool v.0.91b implemented at http://molevol.cmima.csic.es/castresana/Gblocks_server.html and trees developed using FastTree, available at http://www.microbesonline.org/fasttree/ with JTT-CAT models of amino acid evolution, 1,000 tree re-samples and local support values estimated with the Shimodaira-Hasegawa test.

**Table 1.** *Bean yellow mosaic virus* strains/isolates used in comparative sequence analyses, with their corresponding host, origin and GenBank accession numbers

| ID | Host | Strain/isolate | Origin |
| --- | --- | --- | --- |
| AB439729 | *Gladiolus hybrid* cultivar | Gla | Japan |
| KF155419 | *Gladiolus* sp | CK-GL4 | India |
| KM114059 | *Gladiolus dalenii* cv. Sylvia | CK-GL2 | India |
| AB079886 | *Gladiolus* sp | M11 | Japan |
| AB079887 | *Gladiolus* sp | IbG | Japan |
| AB373203 | *Pisum sativum* (pea) | CS | Japan |
| AB439730 | *Gladiolus hybrid* cultivar | G1 | Japan |
| AB439731 | *Vicia faba* | 90-2 | Japan |
| AB439732 | *Trifolium pratense* | 92-1 | Japan |
| AM884180 | *Eustoma russellianum* | lisianthus | Taiwan |
| AY192568 | *Nicotiana benthamiana* | GDD | USA |
| D83749 | *Nicotiana benthamiana* | MB4 | Japan |
| DQ641248 | *Lupinus* sp | W | USA |
| FJ492961 | *Freesia* sp | Fr | Corea del sur |
| FJ492961 | *Freesia* sp | Fr | South Korea |
| HG970847 | *Lupinus cosentinii* | MD1 | Australia |
| HG970850 | *Lupinus cosentinii* | MD7 | Australia |
| HG970851 | *Lupinus angustifolius* | SP1 | Australia |
| HG970858 | *Lupinus angustifolius* | ES55C | Australia |
| HG970863 | *Lupinus angustifolius* | AR87C | Australia |
| HG970866 | *Lupinus pilosus* | LP | Australia |
| HG970867 | *Vicia faba* | FB | Australia |
| HG970868 | *Vicia faba* | LPexFB | Australia |
| JN692500 | *Vicia faba* | Vfaba2 | India |
| JX156423 | *Diuris* sp | SW-3 | Australia |
| JX173278 | *Diuris magnifica* | KP2 | Australia |
| KF155409 | *Gladiolus* cv. Aldebaran | CK-GL1 | India |
| KF155414 | *Gladiolus* cv. Tiger flame | CK-GL | India |
| KF155420 | *Gladiolus* cv. Regency | CK-GL5 | India |
| KT934334 | sunflower | BYSun | Iran |
| U47033 | *Vicia faba* | BYMV-S | Australia |

Reference ID= GenBank accession numbers

## Results and Discussion

Plants showing marked foliar mosaic symptoms, typical of viral infection, were collected during the 2015 growing season, in the green belt of Córdoba city, Argentina. Preparations of symptomatic tissues were mechanically inoculated onto healthy broad bean plants in the greenhouse. All inoculated plants developed symptoms similar to those observed in the field (**Figure 1**). In addition, flexuous filamentous particles, typical of potyviruses were observed under the electronic microscope on dip preparations (**Figure 2**) and all samples were positive when tested by Indirect ELISA with the *anti*-*potyvirus* group monoclonal antibody. To further advance in the characterization of the ELISA positive samples, total RNA was extracted from a symptomatic leaf and submitted to high-throughput sequencing (Miseq-Illumina instrument). After trimming and filtering a total of 935,520 reads were *de novo* assembled into 17,214 contigs, which were subjected to bulk Blastx searches against refseq viral proteins. Seven contigs ranging from 1,085 to 3,527 nt presented significant hits (E-value = 0) to the polyprotein of *Bean yellow mosaic virus* (BYMV, NP_612218.1). Further curation by cycles of iterative mapping of reads and re-assembly (Debat and Bejerman 2019) resulted in a polished, genome sized, virus sequence of 9,497 nt (mean coverage 45.4X, supported by 2,667 reads). The virus sequence presented a 5’-UTR of 196 nt, a single ORF encoding a polyprotein and a 3’-UTR of 130 nt followed by a poly (A) tail. The structural and functional annotation of the PP suggested unequivocally that the obtained sequence (Accession # MK649741) corresponded to a potyvirus (**Figure 3**). The sequence contains a single large open reading frame of 9,171 nt, with a typical potyvirus genomic organization and showing an overall sequence identity of 77.5% with BYMV strain Cs (AB373203). This large ORF encodes a PP of 3,056 aa (347 kDa). PP comparison with the highest identity Blastp hit (BYMV-Cs) suggested that the PP likely contains ten functional proteins in the canonical order 5’-P1-HC-Pro-P3-6K1-CI-6K2-VPg-NIa-Pro-Nib-CP-3’. Each protein presented characteristic size and functional and structural domains of potyviruses. Our results suggest that the detected strain of BYMV is highly divergent, showing highest identities ranging from 64.4 to 95.5% for each mature predicted protein to the closest reported BYMV-Cs (**Figure 3**). Moreover, corresponding PP cleavage sites at the expected nine positions were identified based on MSA alignments with reference potyvirus sequences and mapping confirmed sites available at http://www.dpvweb.net/potycleavage/species.html. Our results indicate that cleavage sites are canonical and differ with BYMV-Cs only at the 6K1/CI and Nia-VPg/Nia-Pro sites (**Figure 3**). In addition, a conserved +2 ribosomal frameshifting signal of the type GAA_AAA_A was identified based on Chung et al. (2008) which supports the generation of a PIPO overlapping ORF between positions 2,887-3,132 nt. The PIPO ORF would generate an 81 aa (8.9 kDa) PIPO protein showing 66.7% identity with the counterpart of BYMV-Cs. Possible recombination events were screened using the RDP4 tool available at http://web.cbio.uct.ac.za/~darren/rdp.html using the complete genome sequences 30 potyviruses with best Blastn hits to BYYV. The software was implemented with default parameters, and p-value cut-off of 0.01. We found no evidence of recombination event with significant hits in more than half of the tools included in RDP4. All in all, these results based on genetic insights, suggested that the detected virus sequence corresponds to a divergent strain of BYMV. To entertain this hypothesis we generated further phylogenetic studies based on MSA of complete PP of refseqs sequences of members of the *Potyviridae* family. The obtained tree clearly showed that the virus clustered within the *Potyvirus* genus (**Figure 4.A**). In addition, local topology of the tree indicates that the detected virus clusters with BYMV (**Figure 4.B**). In this scenario, we tentatively dubbed BYMV strain ARGbb to the assembled virus sequence.

**Figure 1.**
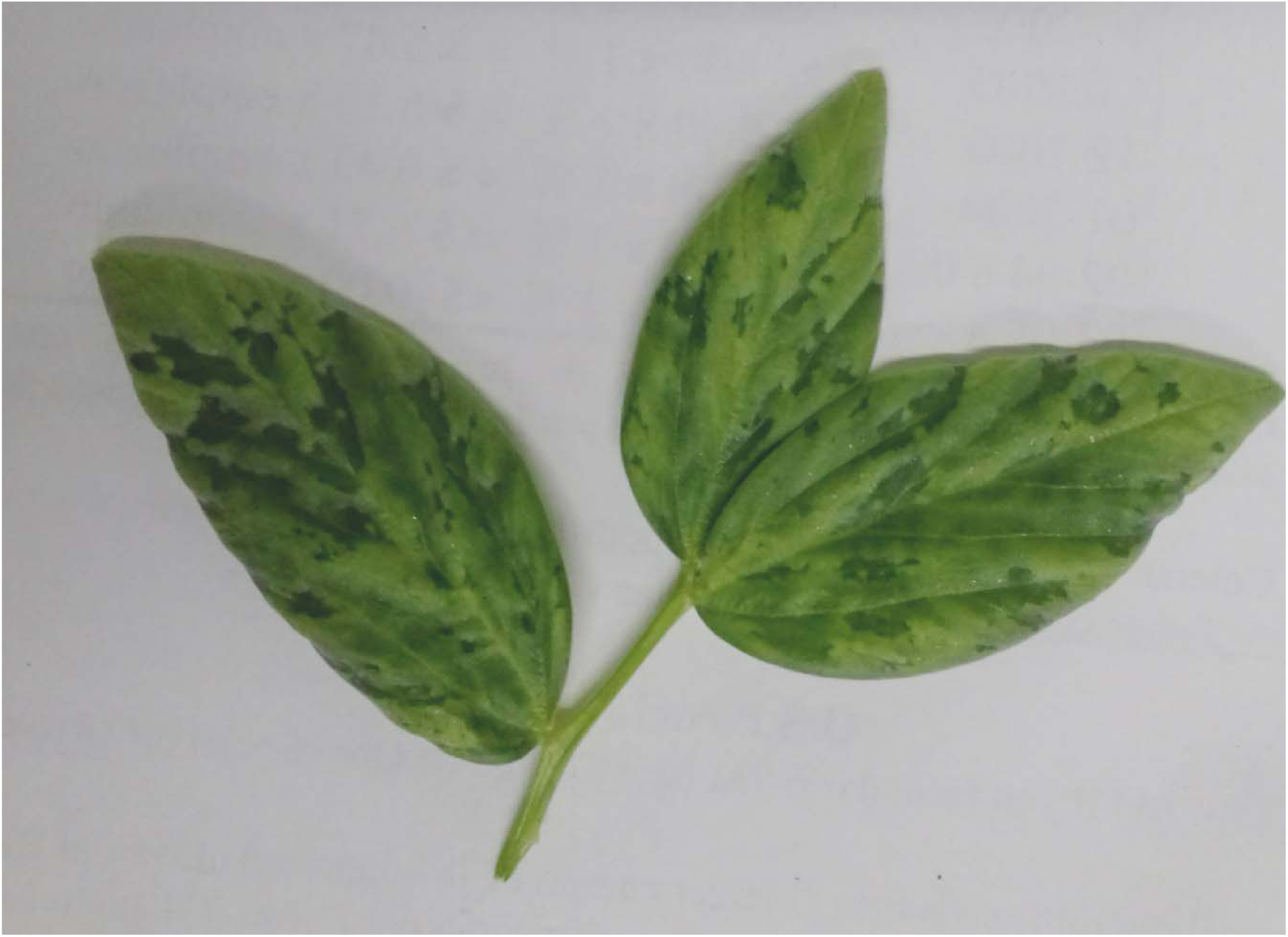
Representative broad bean leaflet showing mosaic symptoms 14 days after inoculation. Samples with mosaic symptoms were mechanically inoculated onto healthy broad bean plants, using sodium phosphate buffer plus silicon carbide. The plants were transferred into a greenhouse and observed daily until symptoms development.

**Figure 2.**
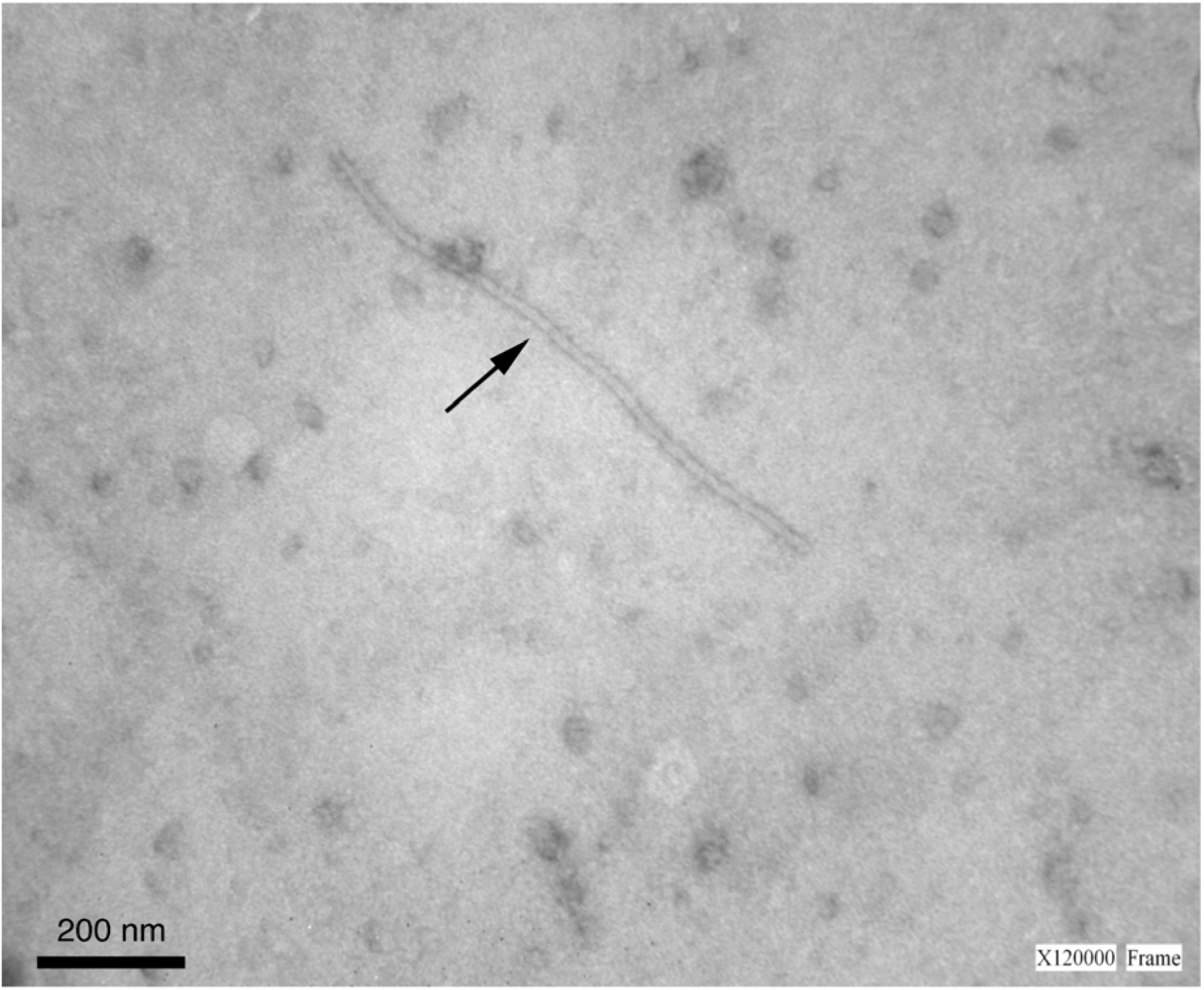
Representative micrograph showing flexuous filamentous particles (black arrow), typical of potyviruses, observed under electronic microscope on dip preparations of symptomatic leaf samples. Scale bar = 200 nm.

**Figure 3.**
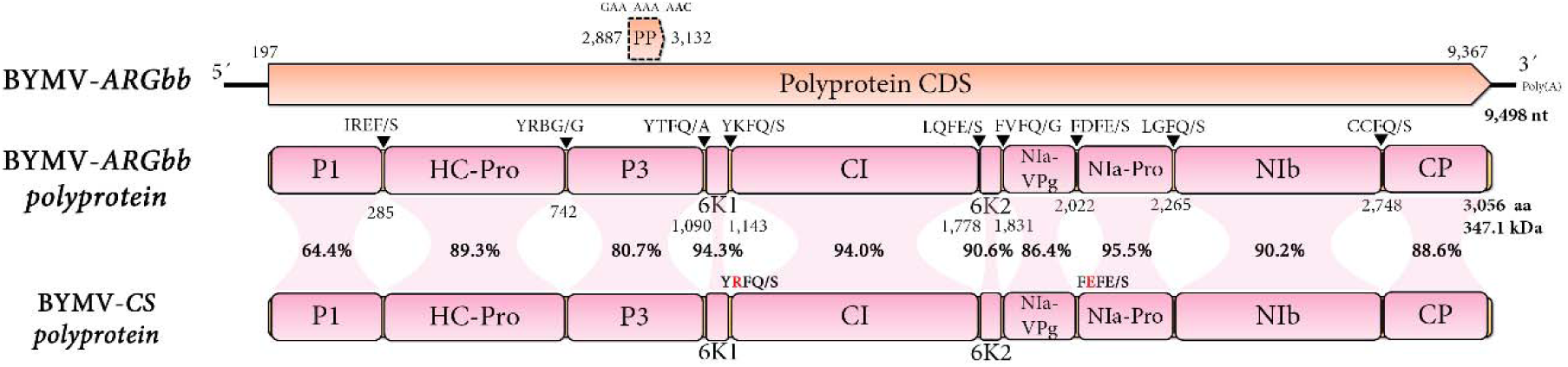
Genome graph depicting architecture and predicted gene products of BYMV-ARGbb. The coding sequences (CDS) are shown in light orange arrow rectangles, start and end coordinates are indicated. Gene products are depicted in curved pink rectangles and aa coordinates are indicated below. Cleavage sites are indicated by black triangles and sequences shown above. Variants of BYMV-Cs are shown in red font. Pairwise aa identity of each protein with that of BYMV-Cs is indicated as %.

**Figure 4.**
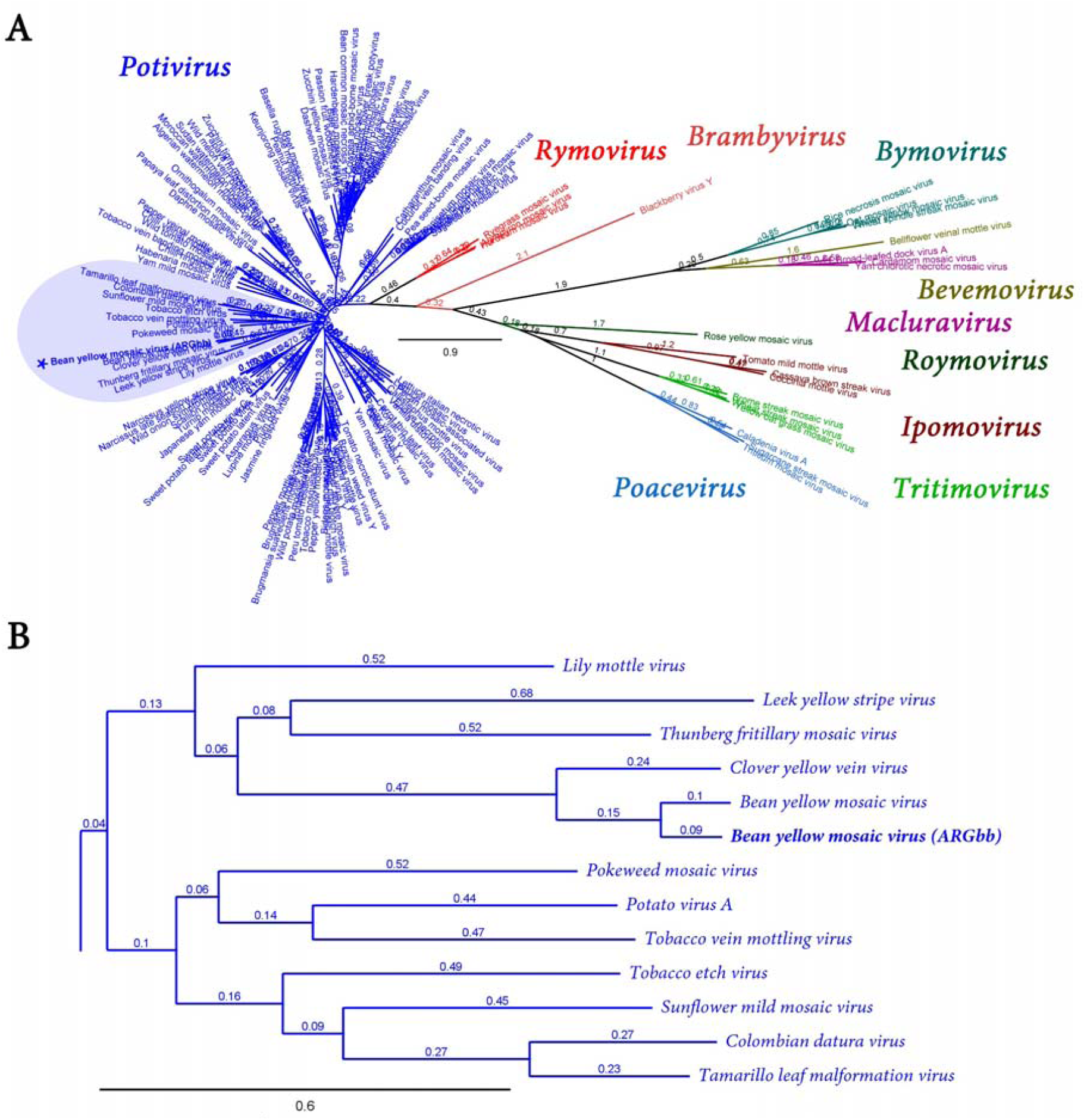
Phylogenetic insights of BYMV-ARGbb. (A) Radial unrooted maximum likelihood phylogenetic tree based on amino acid alignments of the complete polyprotein of BYMV and reference sequences of members of the *Potyviridae* family of viruses. Genera are indicated by color. (B) Magnification or a rooted visualization of the preceding tree showing local topology of the light blue shaded area of A. The scale bar indicates the number of substitutions per site. Node labels indicate FastTree support values.

BYMV is worldwide distributed, has a wide range of natural hosts that includes both monocots and dicots plants, and is the most common cause of mosaic symptoms in faba beans, resulting in yield losses ranging from 81 to 39%, according to the plant growth stage at the time of infection. This virus is transmitted in a non-persistent manner by numerous aphid species and by broad bean seeds, with transmission values between 0.1 and 15% (Berlandier et al. 1997; Kumari and Makkouk 2007; Latham and Jones 2001; Sasaya et al. 1993). It has also been found that BYMV occasionally produces necrotic rings and discoloration in faba bean seeds, diminishing their commercial quality (Kaiser 1973). On the other hand, it has been proven that different isolates of BYMV differ in terms of degree of pathogenicity and serological properties (Barnett et al. 1987; Jones and Diachun 1977; Granett and R. 1975; Cheng and Jones 2000). In general, phylogenetic studies, based on the coat protein sequence of BYMV, established the existence of seven groups, a general one, with a wide hosts range, and six others named according to the original hosts (broad bean, canna, lupine, monocot, pea and W) (Wylie et al. 2008). Further analysis based on the entire virus genome, revealed the presence of nine distinct groups, including the subdivision of the former general group into three new ones (Kehoe et al. 2014), and recently Kaur et al. (2018) suggested that the General group IV of the previous classification could be divided into two subgroups (IVa and IVb). In this scenario, we generated additional phylogenetic analyses to further assess evolutionary relations of the detected virus among BYMV. According to these new grouping, the Argentinean broad bean isolate (BYMV ARGbb), based on entire genome maximum likelihood trees, belongs to group IX together with the Japanese pea isolate (**Figure 5**). Notably, BYMV was previously detected infecting gladiolus and soybean in Argentina (Arneodo et al. 2005; Campos et al. 2014), although only the molecular characterization of the soybean isolate was carried out, and it was shown that this isolate, unlike the broad bean isolate, constituted a new monotypic strain that clustered near the monocot group, evidencing that there are at least two BYMV virus strains present in Argentina. Given the economical importance of this virus and its associated disease, the results presented here are a pivotal first step oriented to explore the eventual incidence and epidemiological parameters of BYMV in broad bean in Argentina.

**Figure 5.**
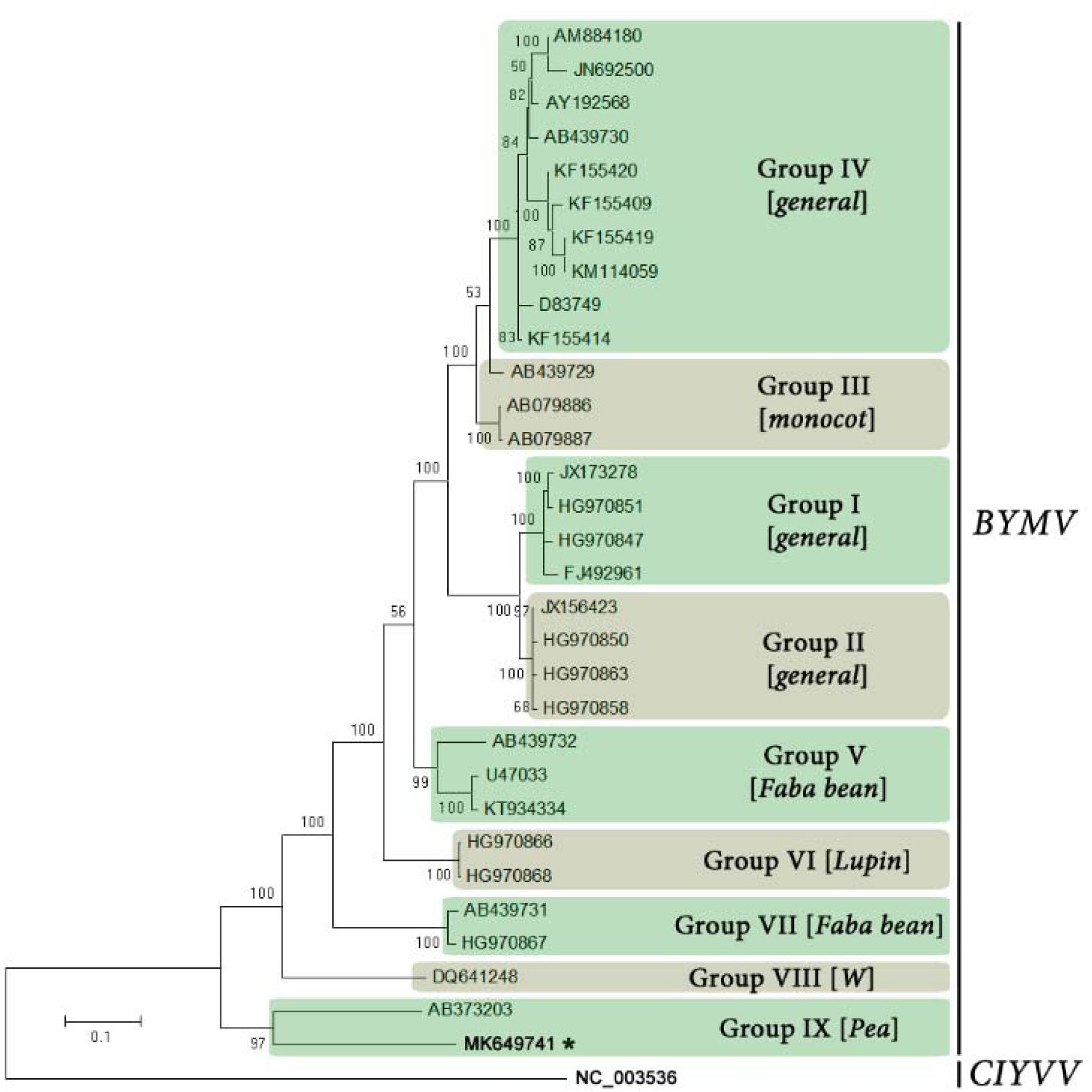
Maximum likelihood tree showing the phylogenetic relationship between Bean yellow mosaic virus isolated from broad bean in Argentina (BYMVFbArg) with other BYMV sequences listed in Table 1. A *Clover yellow vein virus* (ClYVV) isolate was used as the outgroup sequence. The alignment was produced using the complete genome nucleotide sequences. Phylograms were generated with MEGA 5.2 using the Tamura Nei model. Numbers at branches indicate the percentage of 1,000 bootstrap replications where values were above 50%. The evolutionary distance scale is in the units of the number of nucleotide substitutions per site.

## Data availability

The sequence of Bean yellow mosaic virus strain ARGbb has been deposited in NCBI GenBank under accession number MK649741

## Supporting information

Virus sequence MK649741

## Acknowledgements

This study was financed in part by the the Argentine Government, through INTA’s projects PNPV 1135022 and PNHFA 1106075.

## Compliance with ethical standards

### Human and animal rights

This article does not contain any studies with human participants or animals performed by any of the authors.

## References

Arneodo, J. D., de Breuil, S., Lenardon, S. L., Conci, V. C., & Conci, L. R. (2005). Detection of Bean yellow mosaic virus and Cucumber mosaic virus infecting gladiolus in Argentina. Agriscientia, 22(2), 87–89.

Barnett, O. W., Randles, J. W., & Burrows, P. M. (1987). Relationships among Australian and North American isolates of the bean yellow mosaic potyvirus subgroup. Phytopathology, 77, 791–799.

Berlandier, F. A., Thackray, D. J., Jones, R. A. C., Latham, L. J., & CartwrighT, L. (1997). Determining the relative roles of different aphid species as vectors of cucumber mosaic and bean yellow mosaic viruses in lupins. Annals of Applied Biology, 131(2), 297–314, doi:10.1111/j.1744-7348.1997.tb05158.x.

Campos, R. E., Bejerman, N., Nome, C., Laguna, I. G., & Rodríguez Pardina, P. (2014). Bean Yellow Mosaic Virus in Soybean from Argentina. Journal of Phytopathology, 162(5), 322–325, doi:10.1111/jph.12185.

Clark, M. F., & Adams, A. N. (1977). Characteristics of the microplate method of enzyme-linked immunosorbent assay for the detection of plant viruses. Journal of General Virology, 34, 475–483.

Cheng, Y., & Jones, R. A. C. (2000). Biological properties of necrotic and non-necrotic strains of bean yellow mosaic virus in cool season grain legumes. Annals of Applied Biology, 136(3), 215–227, doi:0.1111/j.1744-7348.2000.tb00028.x.

Chung, B. Y., Miller, W. A., Atkins, J. F., & Firth, A. E. (2008). An over-lapping essential gene in the Potyviridae. Proceedings of the National Academy of Sciences, 105, 5897–5902.

Debat, H. J. (2017). An RNA Virome Associated to the Golden Orb-Weaver Spider Nephila clavipes. [Original Research]. Frontiers in Microbiology, 8(2097), doi:10.3389/fmicb.2017.02097.

Debat, H. J., & Bejerman, N. (2019). Novel bird’s-foot trefoil RNA viruses provide insights into a clade of legume-associated enamoviruses and rhabdoviruses. [journal article]. Archives of Virology, doi:10.1007/s00705-019-04193-1.

FAOSTAT (2017). Food and Agriculture Organization of the United Nations. http://www.fao.org/faostat/es/#data/QC.

Goites, E. (2008). Manual de cultivos para la Huerta orgánica familiar. Buenos Aires, Argentina: INTA.

Graham, P. H., & Vance, C. P. (2000). Nitrogen fixation in perspective: an overview of research and extension needs. Field Crops Research, 65(2), 93–106, doi:https://doi.org/10.1016/S0378-4290(99)00080-5.

Granett, A. L., & R., P. (1975). Partial purification and serological relationship of three strains of bean yellow mosaic virus. Annals of Applied Biology, 81(3), 413–415, doi:10.1111/j.1744-7348.1975.tb01658.x.

Jones, R. T., & Diachun, S. (1977). Serologically and biologically distinct bean yellow mosaic virus strains. Phytopathology, 67, 831–838.

Kaiser, W. J. (1973). Biology of Bean Yellow Mosaic and Pea Leaf Roll Viruses affecting Vicia faba in Iran. Journal of Phytopathology, 78(3), 253.

Kaur, C., Raj, R., Srivastava, A., Kumar, S., & Raj, S. K. (2018). Sequence analysis of six full-length bean yellow mosaic virus genomes reveals phylogenetic diversity in India strains, suggesting subdivision of phylogenetic group-IV. [journal article]. Archives of Virology, 163(1), 235–242, doi:10.1007/s00705-017-3609-5.

Kehoe, M. A., Coutts, B. A., Buirchell, B. J., & Jones, R. A. C. (2014). Plant virology and next generation sequencing: Experiences with a Potyvirus. [Article]. PLoS ONE, 9(8), doi:10.1371/journal.pone.0104580.

Kumari, S. G., & Makkouk, K. M. (2007). Virus diseases of faba bean (*Vicia faba* L) in Asia and Africa. Plant viruses, 1(1), 93–105.

Latham, L. J., & Jones, R. A. C. (2001). Incidence of virus infection in experimental plots, commercial crops, and seed stocks of cool season crop legumes. Australian Journal of Agricultural Research, 52(3), 397–413, doi:https://doi.org/10.1071/AR00079.

Makkouk, K. M., Kumari, S. G., Hughes, J. d. A., Muniyappa, V., & Kulkarni, N. K. (2003). Other legumes. In G. Loebenstein, & G. Thottappilly (Eds.), Virus and Virus-like Diseases of Major Crops in Developing Countries (pp. 447–476). Dordrecht: Springer Netherlands.

Mowat, W. P., Dawson, S., &. (1987). Detection and identification of plant viruses by ELISA using crude sap extracts and unfractionated antisera. journal of Virological Methods, 15, 133–247.

Muñoz, N., Liu, A., Kan, L., Li, M.-W., & Lam, H.-M. (2017). Potential Uses of Wild Germplasms of Grain Legumes for Crop Improvement. 18(2), 328, doi:10.3390/ijms18020328.

Paladino, I. R., Sokolowski, A. C., Irigoin, J., Rodriguez, H., Gagey, M. C., Barrios, M. B., et al. (2018). Soil properties evaluation in horticultural farms of Florencio Varela, Buenos Aires, Argentina. [journal article]. Environmental Earth Sciences, 77(11), 411, doi:10.1007/s12665-018-7568-2.

Sasaya, T., Iwasaki, M., & Yamamoto, T. (1993). Seed Transmission of Bean Yellow Mosaic Virus in Broad Bean (*Vicia faba*). Japanese Journal of Phytopathology, 59(5), 559–562, doi:10.3186/jjphytopath.59.559.

Singh, D., Sangle, U. R., Kumar, B., Tripathi, H. S., Singh, K. P., & gupta, A. K. (2012). Integrated disease management of Faba bean (*Vicia faba* L.). In A. K. Singh, & B. P. Bhatt (Eds.), Faba bean (Vicia faba L.): a potential leguminous crop of India (pp. 279–301). Patna. India.

Tamura, K., Peterson, D., Peterson, N., Stecher, G., Nei, M., & Kumar, S. (2011). MEGA5: Molecular Evolutionary Genetics Analysis Using Maximum Likelihood, Evolutionary Distance, and Maximum Parsimony Methods. Molecular Biology and Evolution, 28(10), 2731–2739, doi:10.1093/molbev/msr121.

Wylie, S. J., Coutts, B. A., Jones, M. G. K., & Jones, R. A. C. (2008). Phylogenetic Analysis of Bean yellow mosaic virus Isolates from Four Continents: Relationship Between the Seven Groups Found and Their Hosts and Origins. Plant Disease, 92(12), 1596–1603, doi:10.1094/pdis-92-12-1596.

